# How to tuna fish: constraint, convergence and integration in the neurocranium of pelagiarian fishes

**DOI:** 10.1101/2022.12.19.521082

**Authors:** Andrew Knapp, Gizéh Rangel-de Lázaro, Anjali Goswami, Matt Friedman, Kory M Evans, Sam Giles, Hermione T Beckett, Zerina Johanson

## Abstract

Morphological evolution of the vertebrate skull has been explored across a wide range of tetrapod clades, but teleost fishes, accounting for roughly half of all vertebrate species, have largely been overlooked. Here we present the results of a study investigating three-dimensional morphological evolution across 114 species of Pelagiaria, a morphologically diverse clade of open-ocean teleost fishes that includes tuna and mackerel. Despite showing high shape disparity, the majority of taxa are concentrated in fairly restricted regions of morphospace, with taxa from all families falling into three distinct clusters. Phylogenetic signal in shape data is significant but low (K_mult_ = 0.27, *p* = 0.001) and a single-rate Ornstein-Uhlenbeck model of evolution is supported, revealing convergence in shape within and between families. Shape is significantly correlated with body elongation (R^2^ = 0.35, *p* =0.001), but correlation with size, diet, and habitat depth is weak. Integration of the neurocranium is high, supporting the hypothesis that high integration may promote the evolution of more extreme morphologies. Taken together, these results suggest that shape evolution in the pelagiarian neurocranium is constrained by a number of factors, resulting in the repeated evolution of a restricted range of morphologies.

## Introduction

The open ocean is the largest single habitat on Earth. In contrast to structurally complex biodiverse settings like coral reefs (Roberts et al., 2002; Rabosky et al., 2018), it might seem environmentally homogenous, but the wide range of light levels, temperatures, currents, and food availability creates a surprising diversity of ecological niches that may drive adaptation. Most research on the phenotypic evolution of marine fishes focuses on clades (Larouche et al. 2022; Evans et al. 2022) or assemblages (Claverie and Wainwright, 2014) principally associated with shallow-water, coastal settings, often in proximity to reefs. However, recent surveys of fishes across marine habitats reveal that some settings are unexpected hotspots of phenotypic evolution (Friedman et al. 2019; Martinez et al. 2021). This previous work largely focuses on patterns of morphological diversity or rates of evolutionary change, with an emphasis on extrinsic factors that may-contribute to elevated phenotypic variety (see Collar et al. 2022). Intrinsic aspects potentially facilitating anatomical divergence have, by contrast, been little explored for these groups.

Modularity is an aspect of organismal structure often implicated in mediating patterns of evolutionary diversification. Modularity refers to trait complexes (i.e., ‘modules’) with high internal integration while showing much lower integration between trait complexes (Zelditch and Goswami, 2021). The ability of sets of traits to covary in this way is thought to permit quasi-independent responses to selection. This in turn may result in mosaic evolution over time because modules are able to independently adapt to localised selective pressures and diverge in evolutionary pattern accordingly. Conversely, higher integration between modules is expected to evolve when there is selective pressure for these modules to covary more strongly. Although highly modular structures may be able to evolve a wider range of morphologies, higher levels of integration are thought to promote the evolution of extreme morphologies by partitioning variance along fewer preferred trajectories (Goswami et al., 2014). Modularity has also been hypothesised to increase through evolutionary time to circumvent developmental constraints (Wagner and Altenberg, 1996; Goswami et al., 2014).

Recent studies of integration and modularity have largely focussed on tetrapod clades (Goswami and Polly, 2010; Felice and Goswami, 2017; Randau and Goswami, 2018; Bardua et al., 2019; Watanabe et al., 2019; Felice et al., 2019), but despite accounting for roughly 50% of the 60,000 named species of vertebrates, teleost fishes have received little attention (but see Larouche et al., 2018; Evans et al., 2017a; Evans et al., 2019, Evans et al., 2021). The teleost skull comprises a large number of kinetic and akinetic elements which can be broadly divided into two regions: the rigidly-associated neurocranium, including the skull roof, braincase and upper margin of the orbit (Nelson, 2016); and the splanchnocranium, representing the kinetic elements of the upper and lower jaws, suspensorium, operculum, and other mouth and gill parts (Gregory, 1933). The relatively large number of skull elements in modern teleosts suggests that the structure is highly modular, even in rigid, fused regions such as the neurocranium (Evans et al., 2017b; Evans et al., 2021), though brain development is known to influence neurocranial integration in vertebrates because it signals the development of overlying tissues and spans a number of enclosing elements (Northcutt and Kaas, 1995; Hu and Marcucio, 2009; Evans et al., 2017a). Similarly, the tetrapod skull has been shown to be highly modular despite the loss or fusion of a number of skull elements through evolutionary history (Goswami et al., 2014; Felice and Goswami, 2017; Bardua et al., 2019; Felice et al., 2019; Fabre et al., 2020). The developmental and evolutionary decoupling of kinetic elements may have allowed teleosts to exploit a wide range of niches by enabling the evolution of diverse feeding strategies (Larouche et al., 2022). Understanding patterns of integration is therefore an important step in understanding the evolution of this complex structure in teleosts in context with better-understood tetrapod groups.

Teleosts are separated from tetrapods by some 425 million years of evolution and so establishing homology between skull elements of these groups is difficult (Thomson, 1993; Schultze et al., 2008). Direct comparison is further complicated by the differences in the number and type of skull elements and high kinesis of the mouth and opercular regions of teleosts. The neurocranium, a single, akinetic structure, forms a logical basis for assessing integration and evolution of the teleost skull as an analogue to the rigid tetrapod skull. The neurocranium has four main roles: it protects and supports the brain and cranial sensory organs, provides structure and shape to the skull and anterior body, acts as a rigid anchor to the kinetic splanchnocranial elements, and anchors the spinal column and associated axial musculature important in modulating feeding and respiration (Gregory, 1933; Camp et al. 2015). In contrast to the teleost skull, the tetrapod skull is largely akinetic, incorporating elements involved in feeding (e.g., maxilla, premaxilla) with the neurocranium into a single, rigid structure. Consequently, when treated as a single unit, the tetrapod skull performs additional functions to the teleost neurocranium. Nonetheless, the main roles of the teleost neurocranium, as outlined above, also apply to the cranial region of the tetrapod skull, and this can provide a basis for comparisons between the two groups.

Geometric morphometrics provides a means to capture and statistically quantify morphology across a clade in great detail, allowing for the testing of evolutionary hypotheses in a phylogenetic framework (Goswami et al., 2019). The taxonomic diversity of teleosts poses a challenge to any comprehensive attempt to quantify morphology using this approach because of the difficulty of assembling a representative sample. By focusing on a single clade, it is possible to sample the majority of shape variation and place it in a phylogenetic context, without the need for sampling potentially thousands of individuals. The clade Pelagiaria comprises 286 species of open-ocean fishes in 75 genera and 15 families, including well-known taxa such as tuna (*Thunnus*) and mackerel (*Scomber*) (Miya et al., 2013; Friedman et al., 2019; Arcila et al., 2021). It was only recognised as monophyletic through the application of molecular systematics; traditional morphological classifications dispersed its members between at least six suborders thought to be distantly related to one another. The oldest fossil pelagiarians are Paleocene in age, and paleontologically calibrated molecular phylogenies indicate an origin around the Cretaceous–Paleogene (K/Pg) boundary (Friedman et al. 2019). Fossil evidence shows that familiar modern body plans for at least some of these families had become established no later than the early Eocene (ca. 56 Ma; Bannikov and Monsch, 2011), and these distinctive morphologies may have arisen in as little as 5 - 7 million years (Friedman et al., 2019), suggesting an early burst model of morphological evolution may be supported for this clade. The K/Pg extinction selected against predatory marine fishes (Friedman 2009), suggesting that some pelagiarian lineages exploited these vacated ecological roles in the early Cenozoic. Consequently, this group is ecologically and morphologically diverse, with body shape and diversity of feeding repertoires being notably high for a clade of this size (Friedman et al., 2019). Given the importance of the skull in shaping the anterior body and its role in housing sensory organs and the feeding apparatus, we expect the morphological and ecological diversity of Pelagiaria to be reflected in skull morphology. The relatively small number of taxa and well-resolved phylogeny allows us to conduct a thorough analysis of morphological evolution across the clade.

In this study we conduct a comprehensive analysis of the pelagiarian neurocranium, using three-dimensional geometric morphometrics to determine patterns of morphological disparity, integration, and evolutionary change across the clade.

## Methods

### Data collection, imaging, and landmarking

X-ray micro-computed tomography (μCT) and Computed tomography (for the larger specimens) were used to create high-definition 3D models of the skulls of 114 spirit-preserved pelagiarian specimens, representing 13 of 15 families and 62 of 75 genera (Fig. 1; Supplementary Table S1). The specimens analysed came from repositories at The Natural History Museum (UK), University of Michigan Museum of Zoology (USA), Field Museum of Natural History (USA), Berlin Zoological Museum (Germany), National Museum of Natural History (France), American Museum of Natural History (USA), Australian Museum (Australia), Yale Peabody Museum (USA), and Natural History Museums of Los Angeles County (USA).

**Figure 1:**
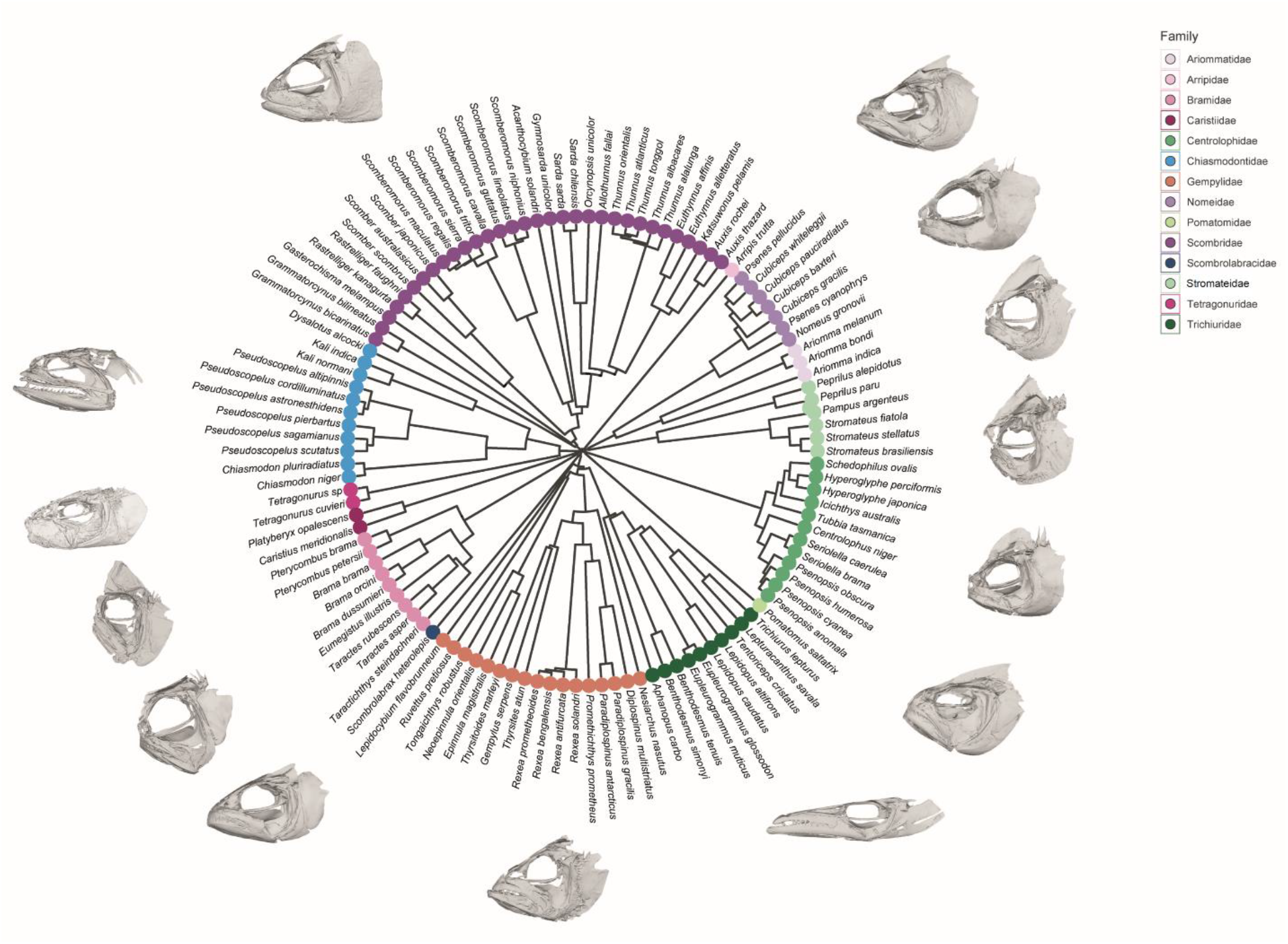
Phylogeny of 114 Pelagiaria species used in this study. Species are coloured according to family. Representative whole-skull meshes are shown for each family. Tree adapted from Friedman et al., 2019 and Arcila et al., 2021.

The specimens were scanned with pixel size and slice thickness ranging between 0.01 and 0.5mm, and an average set of 1657 projections for each individual. The digitised cranial bones were segmented and prepared in Materialise Mimics 21.0 and Geomagic Wrap 2017. The neurocranium of each specimen was segmented out and exported as a ^~^1 million polygon mesh ASCII PLY file.

A landmark scheme was devised to capture as much surface morphology as possible, with 99 anatomical (Bookstein Type I) landmarks to capture homologous points and 102 semilandmark curves to describe sutures and edges (Supplementary Fig. S1). All landmarks and curves were applied with Stratovan Checkpoint (v. 2020.10.13.0859) on the left side of the neurocranium only. Landmarks were exported to R and curves were resampled to standardise the number of semilandmarks, then slid to minimise bending energy. Landmarks were reflected across the midline to improve alignment accuracy (Cardini 2016). A generalised Procrustes alignment (GPA; Rohlf and Slice, 1990) was performed on this set of landmarks (original and reflected), before removing the latter to reduce redundant dimensionality. All subsequent analyses were performed on the aligned, original landmarks.

### Ecological data

Following Friedman et al. (2019), additional information was collected for each taxon. Lateral body elongation was calculated as fineness ratio (standard length/body depth) using the measurement protocol outlined in Martinez et al. (2021). With this method, a circular body has a value of 1 and an elongate body has a value of >>1 (Supplementary Table S1). Following Martinez et al. (2021), each species was assigned to one of three depth zones. These categories are: *Shallow*, equivalent to the epipelagic zone (0-200m, n = 44); *Intermediate*, equivalent to the mesopelagic zone (200-1000m, n = 51); and *Deep*, equivalent to the aphotic bathypelagic and abyssopelagic zones (>1000m, n = 17; Supplementary Table S1). Depth information was gathered from Fishbase (Froese and Pauly, 2022) for all species, based on documented observations. Diet data was binned categorically as large evasive prey (e.g., fishes and squid), gelatinous zooplankton (e.g., jellyfishes, salps), and smaller zooplankton, based on the method used by Friedman et al., 2019 (Supplementary Table S1).

### Phylogeny

We adapted a dated phylogeny for our dataset using results from Friedman et al. (2019), with additional taxa grafted to the tree based on the molecular phylogeny from Arcila et al. (2021; Fig. 1).

### Shape analysis

A principal components analysis (PCA) was performed on the Procrustes-aligned landmark data. Major axes of shape variation were visualised with projected neurocranium shapes, created by warping a mean shape to maximum and minimum PC scores using the ‘tps3d’ function in the R package *Morpho* (R Core Team, 2013; Schlager, 2017). All subsequent analyses were performed in R unless otherwise stated.

Allometry was quantified by regressing Procrustes shape data against centroid size using the ‘procD.lm’ function in *geomorph* (Adams and Otárola-Castillo, 2013). We further quantified allometry in cranial shape after accounting for phylogenetic effects with the ‘procD.pgls’ function in *geomorph*. Phylogenetic signal in shape data was calculated for the whole neurocranium and individual modules using a multivariate adaptation of the *K* statistic, *K_mult_*, implemented with the function ‘physignal’ in *geomorph* (Adams and Otárola-Castillo, 2013).

To assess morphological convergence in neurocranium shape, a k-means cluster analysis was used to identify major clusters within the shape space with the ‘kmeans’ function in R, using the first 35 PC scores (accounting for 95% of total shape disparity). Mean shape was calculated for each cluster using the ‘mshape’ function in *geomorph*, and these shapes were visualised by warping a neurocranium mesh using the ‘tps3d’ function in the R package *Morpho* (Schlager, 2017).

### Modularity analyses

A total of 12 hypotheses of modularity, ranging from a 2-module to a 16-module structure, were developed to encompass a range of developmental, functional, and regional associations of neurocranial elements (see Supplementary material for details). A maximum likelihood approach was implemented with the R package ‘EMMLi’ (Goswami and Finarelli, 2016) to quantify covariance values for all elements across the entire neurocranium. The results of this analysis were used to inform hypotheses 10 – 12, by grouping elements which showed high relative covariance values (Bardua et al., 2019). Each modularity hypothesis was tested with a covariance ratio (CR) analysis using the *geomorph* functions ‘modularity.test’, on raw shape data, and ‘phylo.modularity’, to account for phylogenetic non-independence (Adams and Otárola-Castillo, 2013). The best-supported hypothesis for each subset of the data (raw and phylogenetically corrected shape data) was assessed with the ‘compare.CR’ function in *geomorph*.

### Body shape and ecological data

The correlation of fineness ratio, depth and diet with neurocranium shape was separately assessed with a multivariate regression (for fineness ratio) and a MANOVA (for depth and diet) on the Procrustes shape data, using shape as the dependent variable for each category. These analyses were performed with the R package *geomorph* (Adams and Otárola-Castillo, 2013). The raw shape data were assessed with the ‘procD.lm’ function, and the ‘procD.pgls’ function to account for phylogenetic distance.

Martinez et al. (2021) found that body shape disparity varied significantly with depth in oceanic teleosts, with deep-sea species showing twice the disparity of shallow-water taxa. To test if this was the case in Pelagiaria, we performed a pairwise comparison of shape disparity with depth on our dataset with the function ‘morphol.disparity’ in *geomorph* (Adams and Otárola-Castillo, 2013).

### Evolutionary modelling

We used BayesTraitsV3 (Meade and Pagel, 2014) to estimate evolutionary rates of the entire neurocranium, along with each individual module separately, using the scores of phylogenetic PCs that combined account for >95% of total shape variation. We tested 10 alternative models: fixed and variable rate Brownian motion (BM) and Ornstein-Uhlenbeck (OU), and BM with δ, κ, and λ tree transformations. The best-supported evolutionary model was determined with Bayes Factor in the R package *BTprocessR* (Ferguson-Gow, 2021). Evolutionary rate (σ^2^_mult_) was compared for each module identified in the best-supported modularity hypothesis using the *geomorph* function ‘compare.multi.evol.rates’ (Adams and Otárola-Castillo, 2013).

## Results

### Cranial variation

The results of the PCA for the first 3 PCs are shown in Fig. 2. PC1 accounts for nearly half of shape disparity in the neurocranium (46.1%; Supplementary Fig. S2). It represents a general transition from an elongate neurocranium typical of Gempylidae and Trichiuridae at negative values to the short, deep neurocranium of Stromateidae and Bramidae at positive values. PC2 accounts for 11.6% of total shape disparity and indicates a transition from a wide, dorsoventrally compressed neurocranium with elongate ethmoid, frontal and prefrontal regions at negative PC scores to a laterally compressed, tall neurocranium with large supraoccipital crest and shortened frontal and ethmoid at positive PC scores. Representatives of Scombridae span the entire range of this PC. PC3 accounts for 6.3% of total shape variation and appears to be correlated with relative orbit size, ranging from large orbits seen in Caristiidae at negative PC values to smaller orbits at positive values. The first six PCs cumulatively account for 76% of total shape disparity, indicating that the majority of shape disparity in this dataset is concentrated in a comparatively small number of axes of variation. Phylogenetic signal was significant but low across the whole neurocranium (K_mult_ = 0.27, *p* = 0.001).

**Figure 2:**
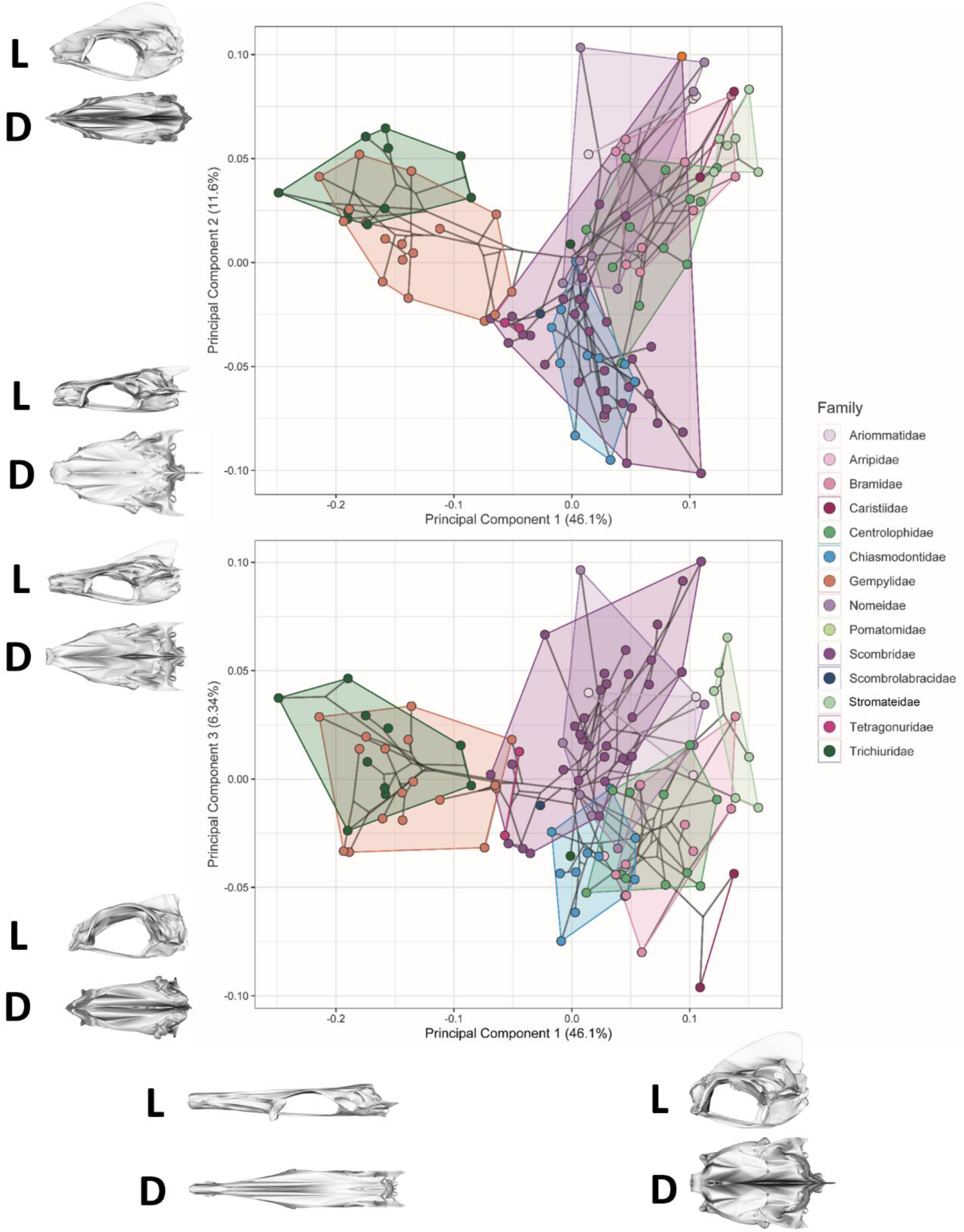
Phylomorphospace of 114 genera of Pelagiaria. Shown are PCs 1 and 2 (top panel) and PCs 1 and 3 (bottom panel). First three PCs cumulatively account for ^~^64% of the total shape disparity. Specimens are coloured according to family, with coloured convex hulls bounding the morphospace ranges of each family. Projected shapes for extreme PC scores are shown as warped meshes as **L:** left lateral and **D:** dorsal view for each PC axis.

Allometry is significantly correlated with neurocranium shape, but with a small effect size, both in raw shape data (R^2^ = 0.04, *p* = 0.002) and after accounting for phylogeny (R^2^ = 0.03, *p* = 0.003), suggesting that size has little influence on shape across the clade.

### Cluster analysis

An examination of the morphospace occupancy density of PCs 1 and 2 suggests several regions of high occupancy (Fig. 3). The cluster analysis performed on the shape data revealed three major clusters that broadly aligned with these regions, corresponding to three distinct morphotypes: morphotype 1 (‘*Brama*’- type), with a shortened dermethmoid region and tall supraoccipital crest; morphotype 2 (‘*Gempylus*’- type), with elongate dermethmoid region and small/no supraoccipital crest; and morphotype 3 (‘*Scomber*’-type), intermediate in shape between morphotypes 1 and 2. Examining this plot also reveals some apparent convergence on these clusters. This is especially obvious at the boundary between morphotype 1 and morphotype 3, with members of Nomeidae, Ariommatidae, Scombridae, Bramidae and Stromateidae all spanning this region.

**Figure 3:**
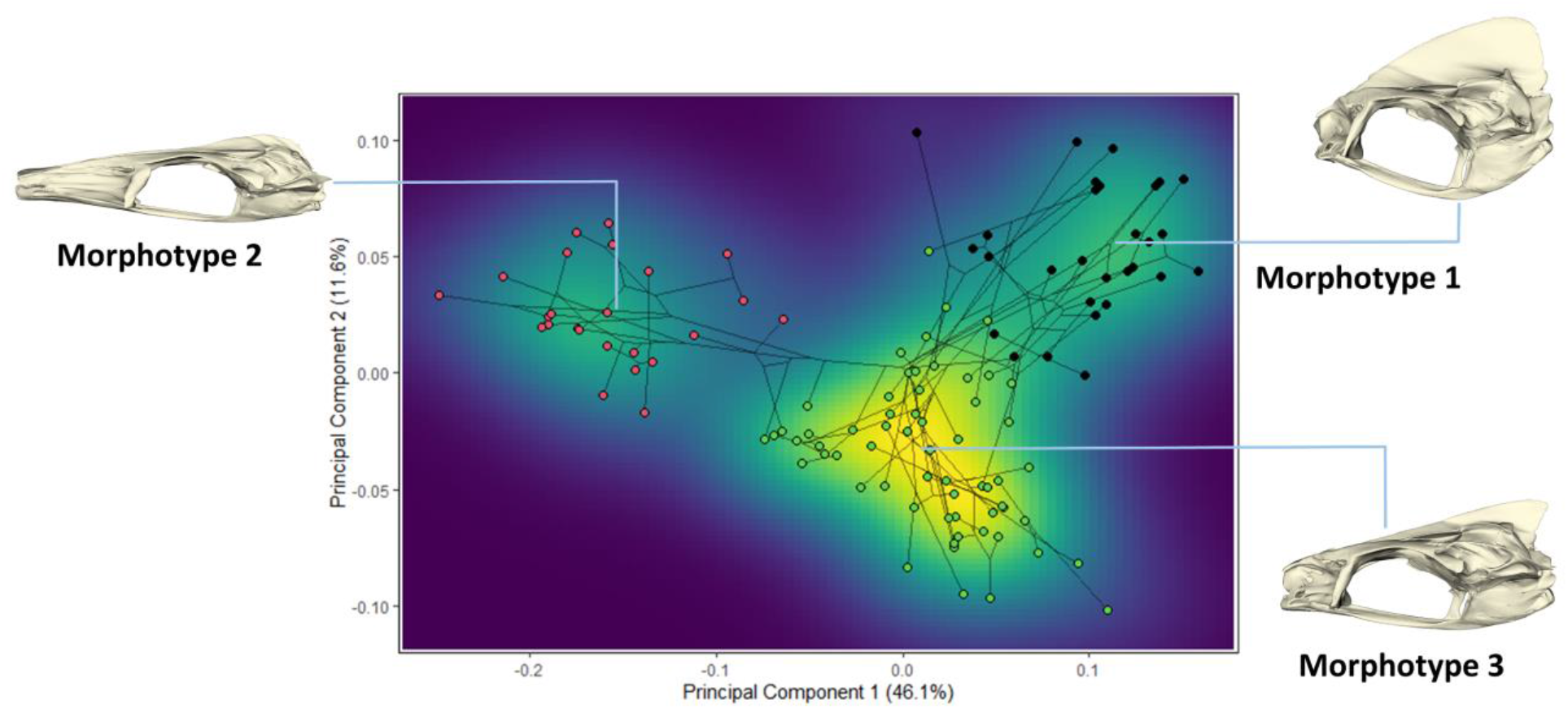
Heatmap showing occupancy density of PCs 1 and 2, with results of cluster analysis of neurocranium morphotypes. Points are coloured according to the morphotype identified from cluster analysis. Meshes represent the mean neurocranium shape for each morphotype.

### Modularity and integration

High integration of the neurocranium is supported by the results of the *compare.CR* analysis, with the strongest support for the 4-module hypothesis (CR = 0.89, *p* = 0.001; Fig. 4). This hypothesis groups dermethmoid, vomer, prefrontal and frontal into the *anterior neurocranium*, the parietal, supraoccipital, exoccipital and basioccipital into the *occipital region*, epiotic, pterotic, opisthotic, sphenotic and prootic into the *otic region*, and the parasphenoid, alisphenoid and basisphenoid into the *sphenoid region*. This result was supported in the compare.CR analysis for both the raw and phylogenetically corrected shape data.

**Figure 4:**
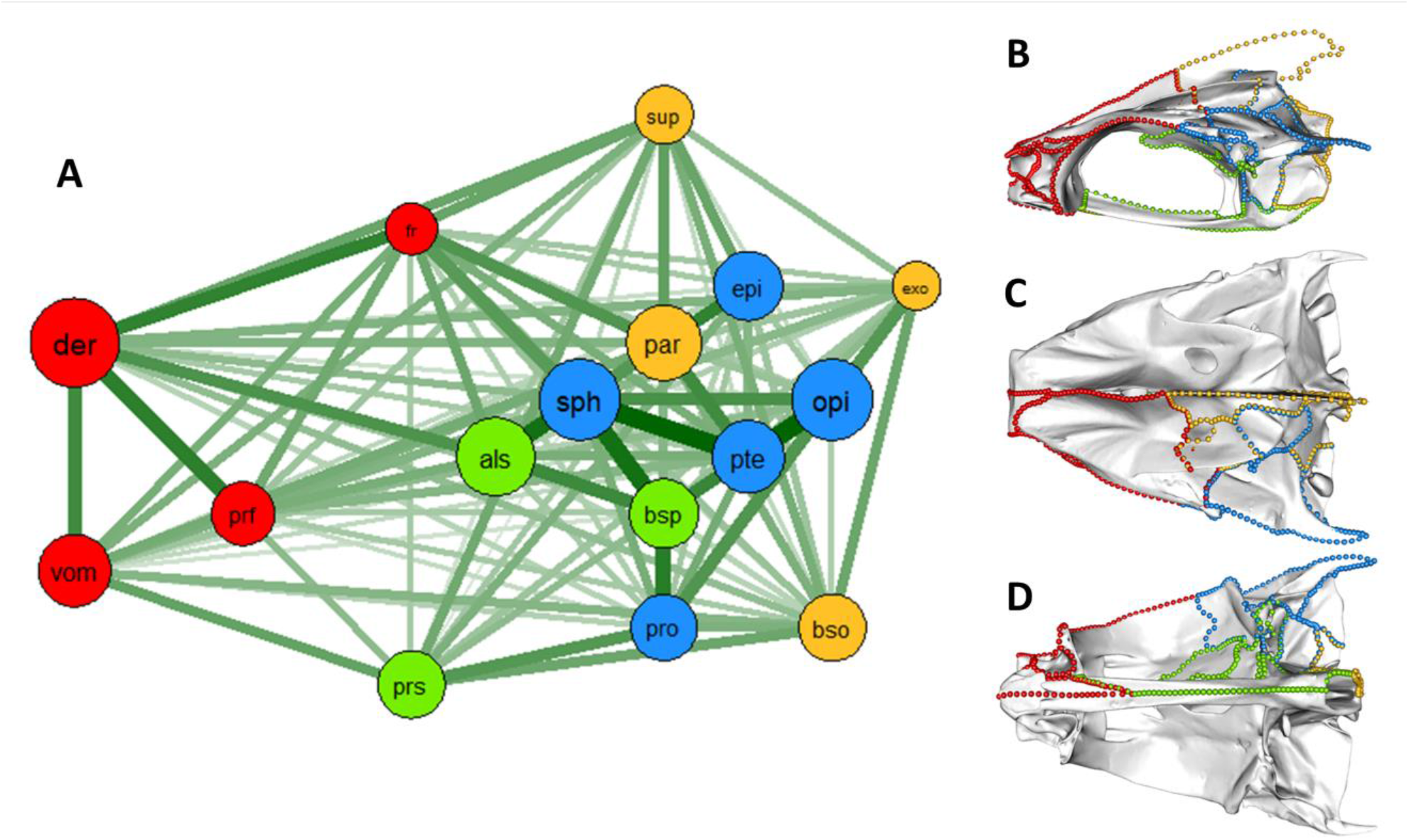
Results of modularity analyses. Network plot (**A**) shows the relative covariance within (circle size) and between (line thickness) skull elements. Meshes show left lateral (**B**), dorsal (**C**) and ventral (**D**) views of neurocranium with landmarks coloured according to module, as per network plot. Abbreviations are **der**: dermethmoid, **prf**: prefrontal, **fr**: frontal, **sup**: supraoccipital, **exo** exoccipital, **epi**: epiotic, **par:** parietal, **sph**: sphenotic, **opi**: opiotic, **pte**: pterotic, **bsp**: basisphenoid, **als**: alisphenoid, **bso**: basioccipital, **pro**: prootic, **prs**: parasphenoid, **vom**: vomer.

### Ecology

Neurocranium shape was weakly correlated with depth (R^2^ = 0.06, *p* = 0.003, Fig. 5), but was non-significant when phylogeny was accounted for (R^2^ = 0.02, *p* = 0.269). Pairwise comparison of depth categories revealed only marginally significant difference in the morphological disparity between the shallow and intermediate groups (Supplementary Table S3), although the deep category had the highest disparity overall (deep = 0.0227, intermediate = 0.0221, shallow = 0.0173). Shape was found to be significantly correlated with fineness ratio (raw shape data: R^2^ = 0.35, *p* = 0.001; phylogenetically corrected: R^2^ = 0.07, *p* = 0.001; Fig. 5), with long, shallow neurocrania being associated with elongate body shapes, and short, deep neurocrania associating with shorter body shapes. There appears to be a general trend in decreasing fineness ratio along PC1, with high fineness ratio values at negative PC scores and low values at positive PC scores (Fig. 5, centre plot). As with depth, dietary categories were significant but with a small effect size in raw shape data (R^2^ = 0.06, *p* = 0.012; Fig. 5, bottom plot) but non-significant after accounting for phylogeny (R^2^ = 0.03, *p* = 0.07).

**Figure 5:**
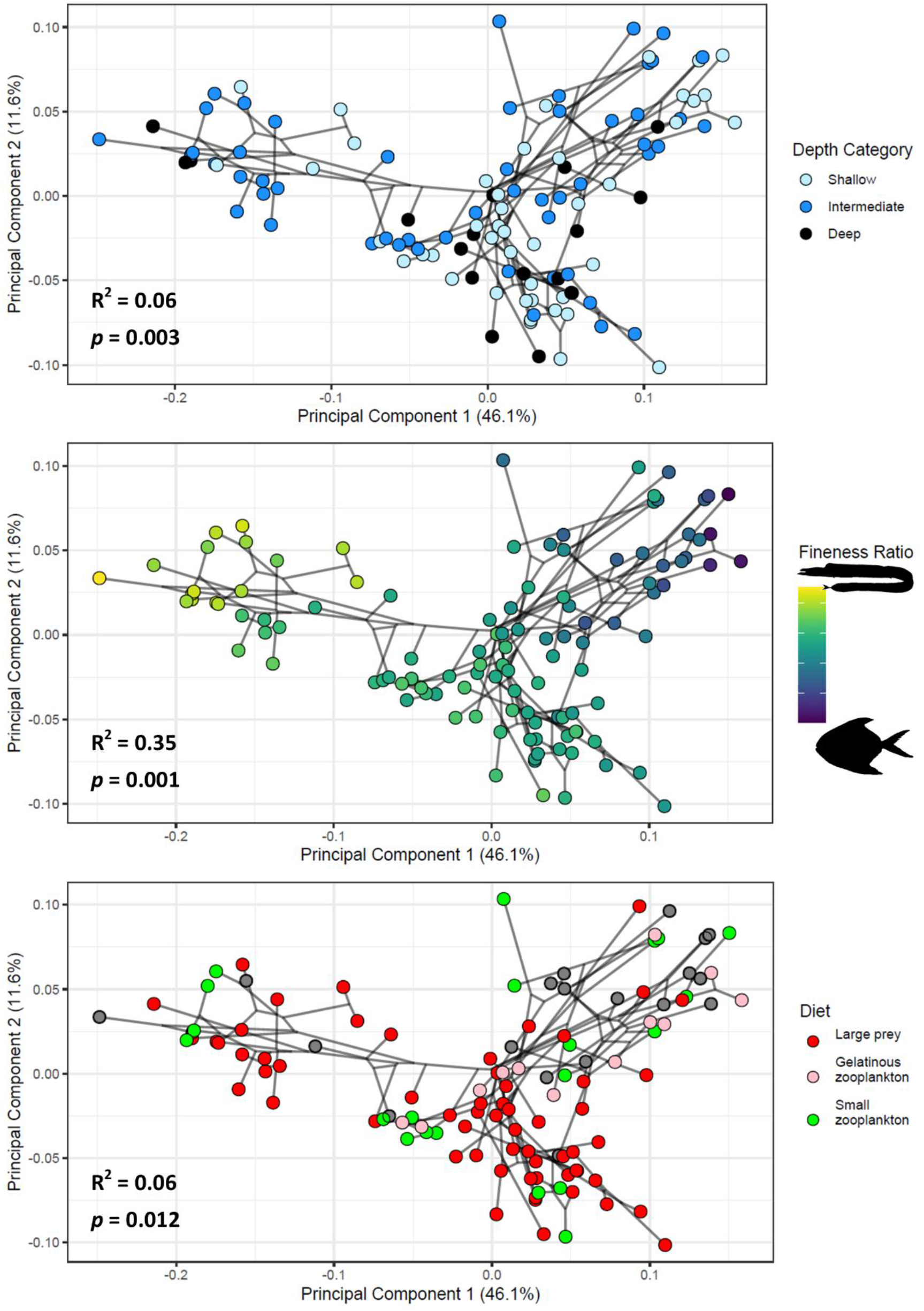
Phylomorphospace of neurocranium shape with specimens coloured by discrete depth (top), continuous fineness ratio (centre), and discrete diet (bottom) values. Effect size (R^2^) and p-values of correlation with raw shape data are shown on each plot.

### Evolutionary modelling

The BayesTraits analysis returned the strongest support for a single-rate OU evolution model for the whole neurocranium, and for each individual module. These results suggest that the pelagiarian neurocranium has undergone consistent rates in shape evolution with little major variation through time but that its shape tends to evolve towards one or more selective optima. Evolutionary rate (**σ^2^_mult_**) was not found to be significantly different between the four modules (Observed rate ratio = 1.96, *p* = 0.316).

## Discussion

Our results reveal that the neurocranium has high shape disparity across Pelagiaria, ranging from long and shallow to short and deep forms. The strong correlation of neurocranium shape with body elongation suggests it plays an important role in structuring and streamlining the anterior body, and it is likely that high integration allows the coordination of evolution and development among skull elements. The grouping of taxa within morphospace suggests that, although highly disparate overall, shape tends to be fairly well-constrained along a small number of axes of variation. This apparent tendency to evolve towards a small number of morphotypes, as suggested by the cluster analysis, is reflected by strong support for a single-rate OU model of evolution. These results support the prediction by Goswami et al. (2014) that high integration can promote the evolution of extreme morphologies by partitioning variance along relatively few trajectories of shape variation.

Phylogenetic integration is expected to play an important role in the evolution and development of complex structures such as the vertebrate skull, because it allows coordinated responses to selection across multiple regions and elements (Klingenberg, 2008; Clune et al., 2013; Goswami et al., 2014; Klingenberg, 2014). Several recent studies have used a geometric morphometric approach to assess evolutionary integration across a range of tetrapod taxa including caecilians (Bardua et al., 2019), birds (Felice and Goswami, 2017; Navalon et al., 2018), archosaurs (Felice et al., 2019; Knapp et al., 2021 Felice at al., 2021), mammals (Goswami and Polly, 2010; Menagaz and Ravosa, 2017; Randau and Goswami, 2018), and squamates (Watanabe et al., 2019). These studies reveal that tetrapod skulls seem to be more modular than those of Pelagiaria. Our present study focussed on the neurocranium, which does not encompass equivalent elements involved in feeding that are found in the tetrapod skull (i.e., maxilla, premaxilla, quadrate, etc.). These other elements might be expected to show more relaxed integration compared to the neurocranium in teleost fishes, particularly given their functional role in feeding. Nonetheless, even the pelagiarian neurocranium on its own seems to show higher integration compared with the equivalent regions of the tetrapod skull, despite incorporating a much higher number of elements. A recent study investigating evolutionary integration in the neurocranium of carangiform fishes also found that integration was high, but linked this to the inclusion of Pleuronectiformes (flatfishes) in the dataset, concluding that the migration of an eye across the sagittal plane of the neurocranium during ontogeny and the resulting directional asymmetry of the neurocrania in this clade was responsible for its high integration (Evans et al., 2021). Although using different landmarking schemes, it is notable that our dataset shows high integration in the neurocranium even in the absence of a morphological feature this extreme.

The significant but weak phylogenetic signal (*K_mult_* = 0.27) recovered from neurocranium shape data suggests a substantial-degree of shape convergence between clades. This is supported by the presence of three distinct clusters within the morphospace (Fig. 3), suggesting that neurocranium shape in Pelagiaria tends to evolve within these somewhat restricted regions of the available morphospace. Gempylidae and Trichiuridae are the only families present in the Morphotype 2 group, but Morphotypes 1 and 3 are populated by taxa from numerous families, and especially obvious crossing-over is apparent between these morphological groups within Nomeidae, Ariommatidae, Scombridae, Bramidae and Stromateidae. It appears that the evolution of certain neurocranium shapes is favoured, with lineages repeatedly evolving towards a restricted range of shape optima, and leaving some regions of morphospace entirely unoccupied (Supplementary Fig. S4). Ornstein-Uhlenbeck evolutionary models describe situations where traits are attracted to selective optima. The strong support found for this model in our study reflects the observed convergence in pelagiarian neurocranial shape (Fig. 4). Furthermore, the single-rate model that was best supported suggests that there have been no major events in the evolutionary history of Pelagiaria that have caused a shift in evolutionary rate, at least in terms of neurocranium shape. For example, the lack of substantial morphological disparity between depth groups in Pelagiaria contrasts with one previous study which showed the deep ocean to be a hotbed of body shape evolution in teleosts (Martinez et al., 2021), so it does not seem that adaptation to different depth zones has notably influenced neurocranial or body shape evolution in this clade. The origination of distinctive crown pelagiarian lineages has been placed near the K/Pg boundary by molecular data (Friedman et al., 2019), and distinctive family-level morphologies are known from Eocene fossils from several localities (Beckett et al., 2018; Friedman et al., 2016), suggesting that morphological disparity was established early in clade diversification. Consequently, an early burst model may be expected to be favoured in Pelagiaria, but this was not supported in our study. Incorporating fossils may affect the best-supported evolutionary model, but the scarcity of well-preserved, three-dimensional neurocranium fossils presently prevents this.

The role of the neurocranium as both a protective structure for the brain and sensory organs, and as a rigid support for the kinetic elements of the skull likely plays an important role in shaping its morphological evolution. The pelagiarian neurocranium does not have an obvious direct mechanical function in the way that, for example, the mandible does (Deakin et al., 2022), and so it is difficult to ascribe an adaptive optimum to any of these shapes based on measurable performance. As in other vertebrates, brain development may influence neurocranium integration in some regions of the pelagiarian neurocranium (Northcutt and Kaas, 1995; Hu and Marcucio, 2009; Evans et al., 2017a). This may account for the high integration and seemingly low disparity of the posterior neurocranium (sphenotic, otic and occipital) when compared with the anterior neurocranium (Figs. 2 and 3). The strong correlation we found between neurocranium shape and body elongation in Pelagiaria highlights its importance in providing structural support and shaping the anterior body. The significant but comparatively weak correlations between neurocranium shape and both diet and depth categories, which disappear after accounting for phylogeny, suggests that these factors are unlikely to play a major role in shaping its morphological evolution. The need for the suspensorium and upper jaw to articulate at several points on the ventral and lateral parts of the neurocranium is likely to place some limits on the relative positions of these contact points to allow for efficient and combined movement of linked kinetic elements (Westneat, 2004). This in turn may depend on the shape and relative position of the elements themselves (Hu et al., 2017). It is possible that the loose association of jaw, suspensorium, and opercular elements in teleosts is a result of relaxed integration across the entire skull. A recently published study incorporating kinetic skull elements suggested that modularity may influence morphological divergence in complex feeding structures in labrid fishes (wrasses) (Larouche et al., 2022). Wrasses have evolved a diverse range of feeding strategies, which are likely to have been driven by the dietary diversity of their reef and shallow water habitats. Although the open ocean is known to be a complex habitat that promotes phenotypic diversity (Friedman et al., 2019; Martinez et al., 2021), pelagiarian taxa do not exhibit certain feeding modes important in wrasses (e.g., grinding and crushing corals, hard-shelled invertebrates, etc.). Consequently, this may have an effect on the phenotypic modularity of the kinetic skull elements in this clade because there is no requirement to evolve crushing jaws and dentition. Incorporating additional skull elements into our dataset will thus let us investigate modularity across the pelagiarian skull and more fully compare across these teleost clades.

## Conclusions

The neurocranium of pelagiarian fishes, a highly diverse clade of open-ocean teleosts, largely follows fundamental hypotheses of the effects of strong integration. The highly disparate shape of the neurocranium across this clade falls along restricted axes of shape variation, and this is likely driven by high covariance between regions of the neurocranium and thus high integration. High integration has previously been predicted to enable the evolution of extreme morphologies, and that appears to be the case in the pelagiarian neurocranium. Neurocranium shape is also strongly correlated with body elongation in Pelagiaria, and species appear to group into three major morphotypes, suggesting some constraints on neurocranium form that result in convergence, and may be a result of this correlation with body shape. Extending this analysis to other skull elements involved in feeding and respiration will allow a more direct comparison with tetrapods, enabling us to establish whether integration remains high across the skull or whether modularity in different regions has allowed the evolutionary and developmental decoupling of these traits to enable diversification in feeding ecologies.

## Supporting information

Supplementary information, including additional tables and figures

## Author contributions

AK, GRL, MF and ZJ conceived the study. AK, GRL, KE, MF, HTB and ZJ collected and processed scan data. AK landmarked scan data. AK and AG analysed data. All authors contributed to writing of the manuscript.

## Acknowledgements

For access to specimens we are grateful to James Maclaine, Ollie Crimmen and Emma Bernard (NHMUK), Peter Rask Möller (NHMD), Peter Bartsch and Edda Assel (MfN), Philippe Béarez and Jonathan Pfliger (MNHN), and Amanda Hay (AMS). For assistance with scanning we thank Vincent Fernandez and Brett Clark (NHMUK), Kristen Mahlow (MfN), Marta Bellato (MNHN), and Jenny Gibson (Royal National Orthopaedic Hospital NHS Trust). We also thank current and past members of the Goswami Lab at the Natural History Museum, London, for valuable feedback and discussion. This study includes data produced in the CTEES facility at University of Michigan, supported by the Department of Earth and Environmental Sciences and College of Literature, Science, and the Arts, and from the Department of Earth Sciences, University of Oxford, Oxford, UK. This study was funded by a Leverhulme Trust grant, no. RPG-2019-113.

