## Supplementary information, including additional tables and figures for "How to tuna fish: constraint, convergence and integration in the neurocranium of pelagiarian fishes"

Andrew Knapp[1*], Gizéh Rangel-de Lázaro[1], Anjali Goswami[1], Matt Friedman[2], Kory M Evans[3], Sam Giles[4], Hermione T Beckett[5], and Zerina Johanson[1]

1. Department of Science, Natural History Museum, London, UK.

2. Department of Earth and Environmental Sciences, University of Michigan, Ann Arbor, USA.

3. Department of Biosciences, Rice University, Houston, Texas, USA.

4. Department of Geography, Earth and Environmental Sciences, University of Birmingham, UK.

5. Department of Biology, King's High School for Girls, Warwick, UK

### Table S1: Information for all specimens used in study.

| Species | Family | Specimen number | Depth group | Diet | Fineness ratio |
|---|---|---|---|---|---|
| *Acanthocybium solandri* | Scombridae | FMNH 44400 | shallow | fish | 6.651 |
| *Allothunnus fallai* | Scombridae | NHMUK 2019.5.24.13 | shallow | fish | 4.588 |
| *Aphanopus carbo* | Trichiuridae | NHMUK 2006.6.27.1 | deep | fish | 16.114 |
| *Ariomma bondi* | Ariommatidae | UMMZ 228950 | intermediate | zooplankton | 3.548 |
| *Ariomma indica* | Ariommatidae | UMMZ 219645 | intermediate | zooplankton | 2.452 |
| *Ariomma melanum* | Ariommatidae | MCZ 168623 6 | intermediate | zooplankton | 4.154 |
| *Arripis trutta* | Arripidae | UMMZ 209265 | shallow | fish | 3.614 |
| *Auxis rochei* | Scombridae | UMMZ 250172 | shallow | zooplankton | 4.829 |
| *Auxis thazard* | Scombridae | UMMZ 238740 | shallow | fish | 4.443 |
| *Benthodesmus simyoni* | Trichiuridae | NHMUK 1972.1.10.64 | intermediate | NA | 33.684 |
| *Benthodesmus tenuis* | Trichiuridae | FMNH 88043 | intermediate | NA | 20.902 |
| *Brama brama* | Bramidae | FMNH 48931 | intermediate | fish | 2.296 |
| *Brama dussumieri* | Bramidae | NHMUK 2004.2.3.107 | shallow | NA | 2.102 |
| *Brama orcini* | Bramidae | MfN 9949 | intermediate | NA | 1.948 |
| *Caristius meridionalis* | Caristiidae | AMS 20071.038.31.002 | intermediate | NA | 1.868 |
| *Centrolophus niger* | Centrolophidae | NHMUK 2014.3.6.7 | deep | fish | 3.911 |
| *Chiasmodon niger* | Chiasmodontidae | UMMZ 228958 | deep | fish | 7.333 |
| *Chiasmodon pluriradiatus* | Chiasmodontidae | NHMUK 1997.5.21.28 | deep | fish | 6.154 |
| *Cubiceps baxteri* | Nomeidae | NHMUK 1997.9.17.27 | shallow | jellyfish | 3.371 |
| *Cubiceps gracilis* | Nomeidae | NHMUK 2019.5.9.256 | shallow | jellyfish | 4.429 |
| *Cubiceps pauciradiatus* | Nomeidae | NHMUK 2004.2.3.99.108 | intermediate | jellyfish | 3.920 |
| *Cubiceps whiteleggi* | Nomeidae | NHMUK 1986.10.6:37-40 | intermediate | jellyfish | 3.368 |
| *Diplospinus multistriatus* | Gempylidae | YPM 0256719 | intermediate | zooplankton | 20.176 |
| *Dysalotus alcocki* | Chiasmodontidae | NHMUK 2017.10.26.237 | deep | fish | 5.900 |
| *Epinnula magistralus* | Gempylidae | UF 233577 | shallow | NA | 4.643 |
| *Eumegistus illustris* | Bramidae | LACM 38573 | intermediate | zooplankton | 2.366 |
| *Eupleurogrammus glossodon* | Trichiuridae | NHMUK 1860.3.19.76 | shallow | fish | 15.810 |
| *Eupleurogrammus muticus* | Trichiuridae | NHMUK 1962_5_4_2 | shallow | fish | 14.286 |
| *Euthynnus affinis* | Scombridae | UMMZ 223134 | shallow | zooplankton | 4.026 |
| *Euthynnus alleteratus* | Scombridae | YPM M39708-71762 | shallow | fish | 3.992 |
| *Gasterochisma melampus* | Scombridae | NHMUK 1868.6.22.2 | intermediate | fish | 3.695 |
| *Gempylus serpens* | Gempylidae | NHMUK 2019.5.9.257 | intermediate | fish | 22.737 |
| *Grammatorcynus bicarinatus* | Scombridae | NHMUK 1872.4.6.25 | shallow | fish | 5.200 |
| *Grammatorcynus bilineatus* | Scombridae | MfN 8003 | shallow | fish | 4.152 |
| *Gymnosarda unicolor* | Scombridae | FMNH 89962 | shallow | fish | 3.890 |

| | | | | | |
|---|---|---|---|---|---|
| *Hyperoglyphe japonica* | Centrolophidae | UMMZ 233106 | deep | zooplankton | 2.900 |
| *Hyperoglyphe perciformis* | Centrolophidae | MNHN IC 1998-1430 | deep | fish | 2.631 |
| *Icichthys australis* | Centrolophidae | NHMUK 2009.5.28.12 | intermediate | NA | 2.778 |
| *Kali indica* | Chiasmodontidae | NHMUK 2018.7.26.214_01 | deep | fish | 4.263 |
| *Kali normani* | Chiasmodontidae | FMNH 88149 | deep | fish | 8.063 |
| *Katsuwonus pelamis* | Scombridae | FMNH 41899 | intermediate | zooplankton | 3.841 |
| *Lepidocybium flavobrunneum* | Gempylidae | MNHN IC 2000.481 | deep | fish | 4.544 |
| *Lepidopus altifrons* | Trichiuridae | FMNH 64192 | intermediate | NA | 12.255 |
| *Lepidopus caudatus* | Trichiuridae | NHMUK 1928.6.24.27 | intermediate | zooplankton | 16.571 |
| *Lepturacanthus savala* | Trichiuridae | UMMZ 219522 | shallow | fish | 15.808 |
| *Neoepinnula orientalis* | Gempylidae | NHMUK 1986.9.8.164 | intermediate | fish | 4.279 |
| *Nesiarchus nasutus* | Gempylidae | FMNH 71422 | deep | fish | 11.797 |
| *Nomeus gronvii* | Nomeidae | NHMUK 1900.11.6.1 | intermediate | jellyfish | 3.625 |
| *Orcynopsis unicolor* | Scombridae | NHMUK 1935.3.5.52 | shallow | fish | 3.764 |
| *Pampus argenteus* | Stromateidae | UMMZ 225567 | shallow | zooplankton | 1.455 |
| *Paradiplospinus antarcticus* | Gempylidae | LACM 10908 | deep | zooplankton | 12.886 |
| *Paradiplospinus gracilis* | Gempylidae | NHMUK 2009.5.18.74 | intermediate | zooplankton | 9.702 |
| *Peprilus alepidotus* | Stromateidae | UMMZ 199053 | shallow | jellyfish | 1.513 |
| *Peprilus paru* | Stromateidae | UMMZ 145907 | shallow | jellyfish | 1.524 |
| *Platyberyx opalescens* | Caristiidae | NHMUK 2005.4.28.1 | deep | NA | 2.045 |
| *Pomatomus saltatrix* | Pomatomidae | UMMZ 111069 | shallow | fish | 3.805 |
| *Promethichthys prometheus* | Gempylidae | UMMZ 250143 | intermediate | fish | 7.667 |
| *Psenes cyanophrys* | Nomeidae | UMMZ 86219 | intermediate | NA | 2.099 |
| *Psenes pellucidus* | Nomeidae | NHMUK 1963.5.14.481 | intermediate | zooplankton | 2.294 |
| *Psenopsis anomala* | Centrolophidae | UMMZ 182977 | intermediate | zooplankton | 2.072 |
| *Psenopsis cyanea* | Centrolophidae | NHMUK 1937.6.28.1 | intermediate | NA | 2.934 |
| *Psenopsis humerosa* | Centrolophidae | AUS I.24816-001 | intermediate | NA | 2.085 |
| *Psenopsis obscura* | Centrolophidae | NHMUK 2002.8.15.5 | intermediate | NA | 3.028 |
| *Pseudoscopelus altipinnis* | Chiasmodontidae | NHMUK 2004.8.17.39 | deep | fish | 5.450 |
| *Pseudoscopelus astronesthidens* | Chiasmodontidae | MCZ 1614171 | deep | fish | 6.453 |
| *Pseudoscopelus cordilluminatus* | Chiasmodontidae | NHMUK 2006.9.19.8 | deep | fish | 4.048 |
| *Pseudoscopelus pierbartus* | Chiasmodontidae | MCZ 164590 | deep | fish | 4.625 |
| *Pseudoscopelus sagamianus* | Chiasmodontidae | NHMUK 1984.1.1.93 | intermediate | fish | 5.886 |
| *Pseudoscopelus scutatus* | Chiasmodontidae | NHMUK 2004.8.17.24 | deep | fish | 6.167 |
| *Pterycombus brama* | Bramidae | UF 168738 | intermediate | NA | 2.103 |
| *Pterycombus petersii* | Bramidae | MNHN 33644 | shallow | NA | 2.700 |
| *Rastrelliger faughni* | Scombridae | NHMUK 1984.1.18.208 | shallow | zooplankton | 4.000 |
| *Rastrelliger kanagurta* | Scombridae | UMMZ 226911 | shallow | zooplankton | 4.032 |
| *Rexea antefurcata* | Gempylidae | NHMUK 1997.5.21.40 | intermediate | fish | 6.040 |
| *Rexea bengalensis* | Gempylidae | NHMUK 1996.9.25.36 | intermediate | fish | 5.602 |
| *Rexea prometheoides* | Gempylidae | FMNH 120779 | intermediate | fish | 5.929 |
| *Rexea solandri* | Gempylidae | UMMZ 216751 | intermediate | fish | 5.090 |
| *Ruvettus pretiosus* | Gempylidae | NHMUK 1938.6.23 | intermediate | fish | 5.043 |
| *Sarda chilensis* | Scombridae | NHMUK 79.10.23.3 | shallow | fish | 4.276 |
| *Sarda sarda* | Scombridae | UMMZ 86218 | shallow | fish | 4.250 |
| *Schedophilus ovalis* | Centrolophidae | NHMUK 2011.2.25.1 | intermediate | jellyfish | 1.892 |
| *Scomber australasicus* | Scombridae | UMMZ 250152 | shallow | zooplankton | 5.325 |
| *Scomber japonicus* | Scombridae | UMMZ 250083 | shallow | zooplankton | 4.736 |
| *Scomber scombrus* | Scombridae | UMMZ 214543 | intermediate | zooplankton | 6.000 |
| *Scomberomorus cavalla* | Scombridae | MfN 1445 | shallow | fish | 4.192 |
| *Scomberomorus guttatus* | Scombridae | MfN 1442 | shallow | fish | 4.766 |
| *Scomberomorus lineolatus* | Scombridae | MNHN IC A 5802 | shallow | fish | 4.247 |
| *Scomberomorus maculatus* | Scombridae | UMMZ 199143 | shallow | fish | 4.527 |
| *Scomberomorus niphonius* | Scombridae | UMMZ 167374 | shallow | fish | 6.461 |
| *Scomberomorus regalis* | Scombridae | UMMZ 143137 | shallow | fish | 5.608 |
| *Scomberomorus sierra* | Scombridae | MNHN IC A 5804 | shallow | fish | 5.489 |
| *Scomberomorus tritor* | Scombridae | MfN 9410 | shallow | fish | 4.274 |
| *Scombrolabrax heterolepis* | Scombrolabracidae | UF 167280 | intermediate | fish | 4.500 |
| *Seriolella brama* | Centrolophidae | NHMUK 1873.12.13.54 | intermediate | jellyfish | 2.833 |
| *Seriolella caerulea* | Centrolophidae | NHMUK 2019.5.24.4 | shallow | jellyfish | 2.109 |
| *Stromateus brasiliensis* | Stromateidae | NHMUK 2019.5.24.1 | shallow | NA | 2.229 |
| *Stromateus fiatola* | Stromateidae | NHMUK 1920.9.7.3 | shallow | fish | 2.356 |
| *Stromateus stellatus* | Stromateidae | NHMUK 1936.8.26.1072 | shallow | NA | 2.463 |
| *Taractes asper* | Bramidae | NHMUK 1953.10.28.2 | shallow | fish | 2.624 |
| *Taractes rubescens* | Bramidae | MCZ 148060 | intermediate | zooplankton | 2.492 |
| *Taractichthys steindachneri* | Bramidae | FMNH 63863 | intermediate | NA | 1.619 |
| *Tentoriceps cristatus* | Trichiuridae | NHMUK 1987.1.23.28 | shallow | fish | 21.429 |

| | | | | | |
|---|---|---|---|---|---|
| *Tetragonurus cuvieri* | Tetragonuridae | NHMD I 39502-001 | intermediate | jellyfish | 6.246 |
| *Tetragonurus sp* | Tetragonuridae | NHMUK 1998.8.9.9520 | intermediate | jellyfish | 6.863 |
| *Thunnus alalunga* | Scombridae | NHMUK 2004.11.1.299 | intermediate | fish | 3.540 |
| *Thunnus albacares* | Scombridae | NHMUK 1973.6.6.1 | shallow | fish | 3.377 |
| *Thunnus atlanticus* | Scombridae | NHMUK 2012.8.15.13 | intermediate | fish | 3.392 |
| *Thunnus orientalis* | Scombridae | FMNH 58745 | intermediate | fish | 3.359 |
| *Thunnus tonggol* | Scombridae | UMMZ 225078 | intermediate | fish | 4.408 |
| *Thyrsites atun* | Gempylidae | NHMUK 1927.12.6.75 | intermediate | fish | 7.939 |
| *Thyrsitoides marleyi* | Gempylidae | NHMUK 1986.9.8.147 | intermediate | fish | 8.660 |
| *Tongaichthys robustus* | Gempylidae | MNHN-IC-2000-0460 | intermediate | NA | 4.278 |
| *Trichiurus lepturus* | Trichiuridae | UMMZ 219710 | intermediate | fish | 13.861 |
| *Tubbia tasmanica* | Centrolophidae | AMS IB.1148 | intermediate | NA | 3.160 |

**Abbreviations: AMS:** Australian Museum, Sydney; **FMNH:** Field Museum of Natural History; **LACM:** Los Angeles County Museum; **MCZ:** Museum of Comparative Zoology, Harvard University; **MfN:** Museum für Naturkunde, Berlin; **MNHN:** Muséum National d'Histoire Naturelle, Paris; **NHMUK:** Natural History Museum, London; **UF:** Florida Museum of Natural History; **UMMZ:** University of Michigan Museum of Zoology; **YPM:** Yale Peabody Museum.

**List of anatomical landmarks**

**Dermethmoid**

1. Forward point of dermethmoid on midline
2. Rear point of dermethmoid on midline at junction with frontals
3. Lateral point of dermethmoid at junction with frontal
4. Forwardmost point of dermethmoid contact with nasal
5. Rearmost junction of dermethmoid with underside of frontal
6. Rearmost lower junction of dermethmoid with prefrontal

**Frontal**

7. Forward junction of frontal and dermethmoid on midline
8. Junction of frontal and supraoccipital on midline
9. Junction of frontal and supraoccipital on margin of dorsal fenestra
10. Junction of frontal and parietal on margin of dorsal fenestra
11. Junction of frontal and parietal on upper lateral ridge
12. Junction of frontal, parietal and pterotic
13. Junction of frontal and pterotic on lower lateral ridge
14. Junction of frontal and
15. Lateral junction of frontal and dermethmoid

**Supraoccipital**

16. Junction of supraoccipital and frontal on midline
17. Junction of supraoccipital and exoccipital on midline
18. Junction of supraoccipital and exoccipital at margin of upper rear fenestra
19. Junction of supraoccipital and epiotic at margin of upper rear fenestra
20. Junction of supraoccipital, epiotic and parietal
21. Junction of supraoccipital and parietal at margin of dorsal fenestra
22. Junction of supraoccipital and frontal at margin of dorsal fenestra

**Exoccipital**

23. Junction of exoccipital and supraoccipital on midline
24. Rear symphysis of exoccipital at upper margin of foramen magnum
25. Upper point of exoccipital symphysis at occipital condyle
26. Lower point of exoccipital symphysis at occipital condyle
27. Forward junction of exoccipital and basioccipital
28. Junction of exoccipital and opisthotic on underside of neurocranium
29. Junction of exoccipital and opisthotic on rear edge of neurocranium
30. Junction of exoccipital and opisthotic on upper surface of neurocranium
31. Outer junction of exoccipital and epiotic
32. Inner junction of exoccipital and epiotic on margin of upper rear fenestra
33. Junction of exoccipital and supraoccipital on margin of upper rear fenestra

**Epiotic**

34. Junction of epiotic, supraoccipital and parietal
35. Junction of epiotic and parietal at margin of lateral fenestra
36. Junction of epiotic, pterotic and exoccipital

37. Junction of epiotic and exoccipital at margin of upper rear fenestra
38. Junction of epiotic and supraoccipital at margin of upper rear fenestra

**Parietal**

39. Junction of parietal, epiotic and supraoccipital
40. Junction of parietal and supraoccipital on margin of dorsal fenestra
41. Junction of parietal and frontal on margin of dorsal fenestra
42. Junction of parietal, frontal and pterotic
43. Junction of parietal and pterotic on margin of lateral fenestra
44. Junction of parietal and epiotic on margin of lateral fenestra

**Opisthotic**

45. Junction of opisthotic and exoccipital on rear edge of neurocranium
46. Junction of opisthotic, pterotic and exoccipital on upper side of neurocranium
47. Junction of opisthotic and pterotic on rear edge of neurocranium
48. Junction of opisthotic, pterotic and prootic on underside of neurocranium
49. Junction of opisthotic and exoccipital on underside of neurocranium

**Pterotic**

50. Junction of pterotic and opisthotic on rear edge of neurocranium
51. Junction between pterotic, exoccipital and opisthotic on upperside of neurocranium
52. Junction of pterotic, exoccipital and epiotic
53. Junction of pterotic and parietal at edge of lateral fenestra
54. Junction of pterotic, parietal and frontal
55. Junction of pterotic and frontal on lower lateral ridge
56. Junction of pterotic and sphenotic at margin of hyomandibular socket
57. Junction of pterotic and basisphenoid at margin of hyomandibular socket
58. Junction of pterotic, opisthotic and prootic
59. Tip of pterotic process

**Basioccipital**

60. Upper point of vertebral centrum contact on midline
61. Lower point of vertebral centrum contact on midline
62. Rear contact point of basioccipital and parasphenoid
63. Forward contact of basioccipital, parasphenoid and prootic
64. Contact of basioccipital, prootic and exoccipital

**Sphenotic**

65. Junction of sphenotic and frontal on orbit margin
66. Junction of sphenotic, frontal and alisphenoid
67. Junction of sphenotic, alisphenoid and basisphenoid
68. Junction of sphenotic and pterotic on margin of hyomandibular socket
69. Junction of sphenotic and pterotic on lower lateral ridge
70. Rear point of lateral sphenotic process

**Alisphenoid**

71. Junction of alisphenoid and frontal

72. Junction of alisphenoid, frontal and sphenotic
73. Junction of alisphenoid, sphenotic and basisphenoid
74. Junction of alisphenoid and basisphenoid

**Prootic**

75. Junction of prootic, parasphenoid and basioccipital
76. Forward junction of prootic and parasphenoid
77. Forward junction of prootic and basisphenoid
78. Junction of prootic, basisphenoid and pterotic

**Basisphenoid**

79. Junction of basisphenoid and alisphenoid
80. Junction of basisphenoid, alisphenoid and sphenotic
81. Junction of basisphenoid and sphenotic on margin of hyomandibular socket
82. Junction of basisphenoid and pterotic on margin of hyomandibular socket
83. Junction of basisphenoid, prootic and pterotic
84. Junction of basisphenoid and prootic at rear of orbital cavity
85. Junction of basisphenoid and basisphenoid process

**Parasphenoid**

86. Rearmost point of parasphenoid on midline
87. Rear junction of parasphenoid and vomer on midline
88. Forward lateral junction of parasphenoid and vomer
89. Point on upper midline of parasphenoid in line with rear edge of prefrontals
90. Contact point of parasphenoid with basisphenoid process on midline
91. Apex of parasphenoid curve at orbit margin
92. Junction between parasphenoid and basisphenoid at rear of orbital cavity

**Vomer**

93. Rearmost junction of vomer and parasphenoid on midline
94. Forward junction of vomer and dermethmoid on midline
95. Lateral junction of vomer and parasphenoid

**Prefrontal**

96. Junction of parethmoid and frontal on orbit margin
97. Tip of parethmoid at articulation with lachrymal
98. Forward junction of parethmoid and vomer
99. Forward upper point of parethmoid at contact with frontal

**List of semilandmark curves (following sutures unless otherwise stated)**

**Dermethmoid**

1. Pts 1 – 2 along midline
2. Pts 2 – 3
3. Pts 3 – 4
4. Pts 3 – 5

5. Pts 5 – 6
6. Pts 6 – 1

**Frontal**

7. Pts 7 – 8 along midline suture
8. Pts 8 – 9
9. Pts 9 – 10
10. Pts 10 – 11
11. Pts 11 – 12
12. Pts 12 – 13
13. Pts 13 – 14
14. Pts 14 – 15 along orbit margin
15. Pts 15 – 7

**Supraoccipital**

16. Pts 16 – 17 along supraoccipital crest midline
17. Pts 17 – 18
18. Pts 18 – 19
19. Pts 19 – 20
20. Pts 20 – 21
21. Pts 21 – 22
22. Pts 22 – 16

**Exoccipital**

23. Pts 23 – 24
24. Pts 24 – 25
25. Pts 25 – 26 along midline of occipital condyle
26. Pts 25 – 26 around perimeter of occipital condyle
27. Pts 26 – 28
28. Pts 27 – 28
29. Pts 28 – 29
30. Pts 29 – 30
31. Pts 30 – 31
32. Pts 31 – 32
33. Pts 32 – 33
34. Pts 33 – 23

**Epiotic**

35. Pts 34 – 35
36. Pts 35 – 36
37. Pts 36 – 37
38. Pts 37 – 38
39. Pts 38 – 34

**Parietal**

40. Pts 39 – 40
41. Pts 40 – 41

42. Pts 41 – 42
43. Pts 42 – 43
44. Pts 43 – 44
45. Pts 44 – 39

**Opisthotic**

46. Pts 45 – 46
47. Pts 46 – 47
48. Pts 47 – 48
49. Pts 48 – 49
50. Pts 49 – 45

**Pterotic**

51. Pts 50 – 51
52. Pts 51 – 52
53. Pts 52 – 53
54. Pts 53 – 54
55. Pts 54 – 55
56. Pts 55 – 56
57. Pts 56 – 57
58. Pts 57 – 58
59. Pts 58 – 50
60. Pts 50 – 59 along rear edge of pterotic
61. Pts 59 – 55 along lower lateral ridge

**Basioccipital**

62. Pts 60 – 61 around perimeter of vertebral centrum contact
63. Pts 61 – 62
64. Pts 62 – 63
65. Pts 63 – 64
66. Pts 64 – 60

**Sphenotic**

67. Pts 65 – 66
68. Pts 66 – 67
69. Pts 67 – 68
70. Pts 68 – 69
71. Pts 69 – 65
72. Pts 65 – 70 along lateral edge of sphenotic process
73. Pts 70 – 68 along rear edge of sphenotic process

**Alisphenoid**

74. Pts 71 – 72
75. Pts 72 – 73
76. Pts 73 – 74
77. Pts 74 – 75

**Prootic**

78. Pts 75 – 76
79. Pts 76 – 77
80. Pts 77 – 78
81. Pts 78 – 75

**Basisphenoid**

82. Pts 79 – 80
83. Pts 80 – 81
84. Pts 81 – 82
85. Pts 82 – 83
86. Pts 83 – 84
87. Pts 84 – 85
88. Pts 85 – 79

**Parasphenoid**

89. Pts 86 – 87
90. Pts 87 – 88
91. Pts 88 – 89
92. Pts 89 – 90
93. Pts 90 – 91
94. Pts 91 – 92
95. Pts 92 – 86

**Vomer**

96. Pts 93 – 94
97. Pts 94 – 95
98. Pts 95 – 93

**Prefrontal**

99. Pts 96 – 97 along lateral edge of prefrontal
100. Pts 97 – 98
101. Pts 98 – 99
102. Pts 99 - 96

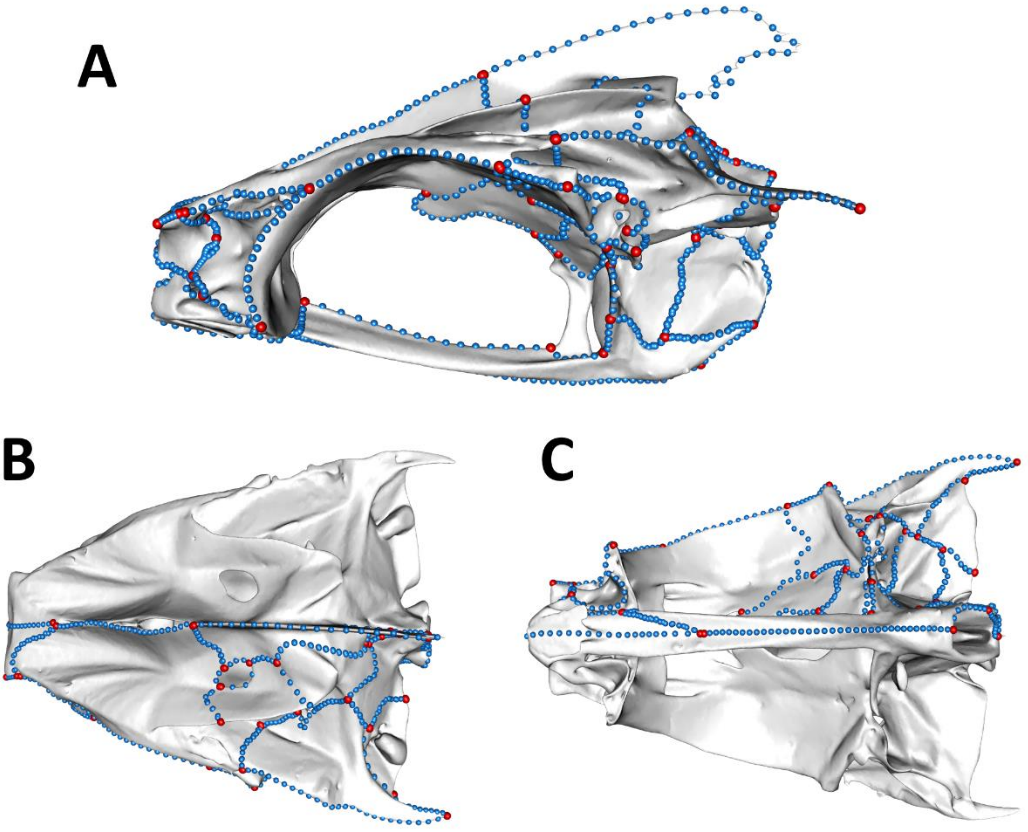


**Figure S1: Landmark layout for Pelagiaria neurocranium.** Shown on a mesh of *Thunnus tonggol* are anatomical landmarks (red points) and semilandmark curves (blue points). Refer to landmark guide for descriptions of placement. **A:** left lateral view; **B:** dorsal view; **C:** ventral view.

**Neurocranium elements (as numbered in modularity hypotheses, below)**

1. Dermethmoid
2. Frontal
3. Supraoccipital
4. Exoccipital
5. Epiotic
6. Parietal
7. Opisthotic
8. Pterotic
9. Basioccipital
10. Sphenotic
11. Alisphenoid
12. Prootic
13. Basisphenoid
14. Parasphenoid
15. Vomer
16. Prefrontal

**Modularity hypotheses**

A maximum likelihood approach, EMMLi (Goswami and Finarelli, 2016) was used to quantify intra- and inter-element covariance values, the results of which were used to inform hypotheses 10 – 12. These hypotheses grouped regions with relatively high levels of between-element integration (see Bardua et al., 2019)

1. All bones separate modules.
2. Anterior/posterior modules: (1(1,2,14,15,16), 2(3:13)).
3. Anterior/posterior modules, parasphenoid part of posterior: (1(1,2,15,16), 2(3:14)).
4. Anterior/posterior modules, separate supraoccipital: (1(1,2,14,15,16), 2(3), 3(4:13)).
5. Anterior/posterior modules, parasphenoid part of posterior, separate supraoccipital: (1(1,2,15,16), 2(3), 3(4:14)).
6. Anterior region, otics, sphenoids and occipitals grouped: (1(1,2,15,16), 2(3,4,6,9), 3(5,7,8,10,12), 4(11,13,14)).
7. Embryology hypothesis I (Kague et al. 2012), Mesoderm and Neural Crest origins, frontal as neural crest: (1(1,2,8,10,11,15,16),2(3,4,5,6,7,12,13,14)).
8. Embryology hypothesis II (Kague et al. 2012), Mesoderm and Neural Crest origins, frontal as mesoderm: (1(1,8,10,11,15,16),2(2,3,4,5,6,7,12,13,14)).
9. Same as embryology hypothesis II (Kague et al. 2012), Mesoderm and Neural Crest origins, frontal as mesoderm, separate supraoccipital: (1(1,8,10,11,15,16), 2(2,4,5,6,7,12,13,14), 3(3))
10. Based on EMMLi results, modules are merged if between-module value is <0.1: (1(1,2,6), 2(3), 3(4,7,8,10), 4(5), 6(9), 7(11), 8(12), 9(13), 10(14), 11(15), 12(16)).
11. Based on EMMLi results, modules are merged if between-module value is <0.15: (1(1,2,3,4,6,7,8,10,12,13,16), 2(5), 3(9), 4(11), 5(14), 6(15)).
12. Based on EMMLi results, modules are merged if between-module value is <0.1 or above overall lowest internal value: (1(1,2,6,16), 2(3), 3(4,7,8,10,11,12,13), 4(5), 5(9), 6(14), 7(15)).

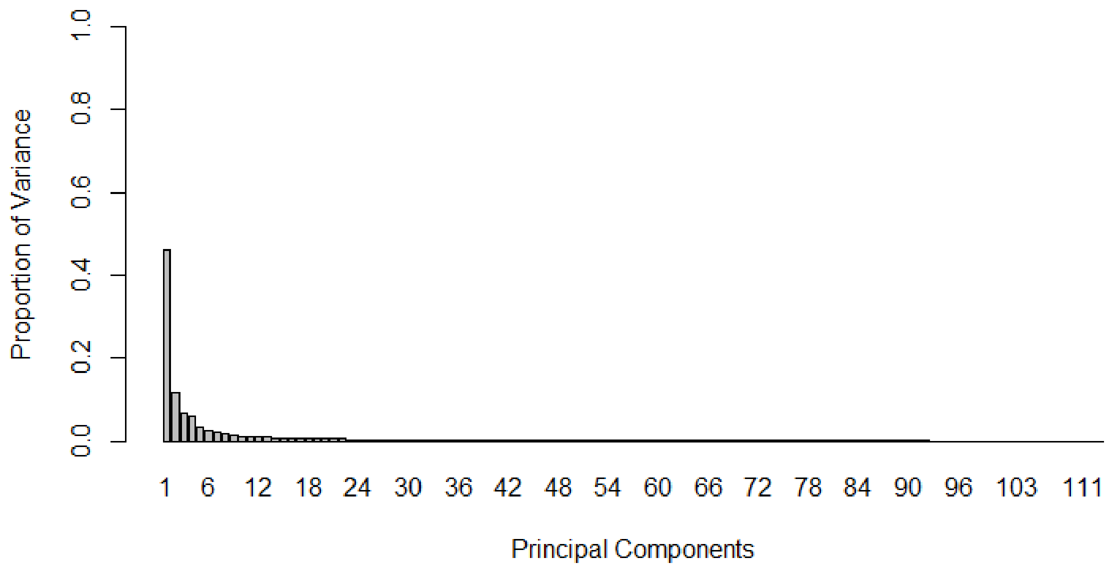

**Figure S2:** Scree plot of PC contributions to shape variation

**Table S2: Phylogenetic signal for whole neurocranium and separate modules.**

| | Phylogenetic signal ($K_{mult}$) | *p* | Evolutionary rate ($\sigma^2_{mult}$) |
|---|---|---|---|
| **Anterior neurocranium** | 0.329 | 0.001 | 1.46 |
| **Occipital** | 0.261 | 0.001 | 1.25 |
| **Otic region** | 0.246 | 0.001 | 0.75 |
| **Sphenoid** | 0.222 | 0.001 | 0.84 |
| **Whole neurocranium** | 0.27 | 0.001 | n/a |

**Table S3: Pairwise p – values for differences in shape disparity for depth categories**

| | Deep | Intermediate | Shallow |
|---|---|---|---|
| **Deep** | 1.0 | 0.89 | 0.12 |
| **Intermediate** | 0.89 | 1.0 | 0.05 |
| **Shallow** | 0.12 | 0.05 | 1.0 |

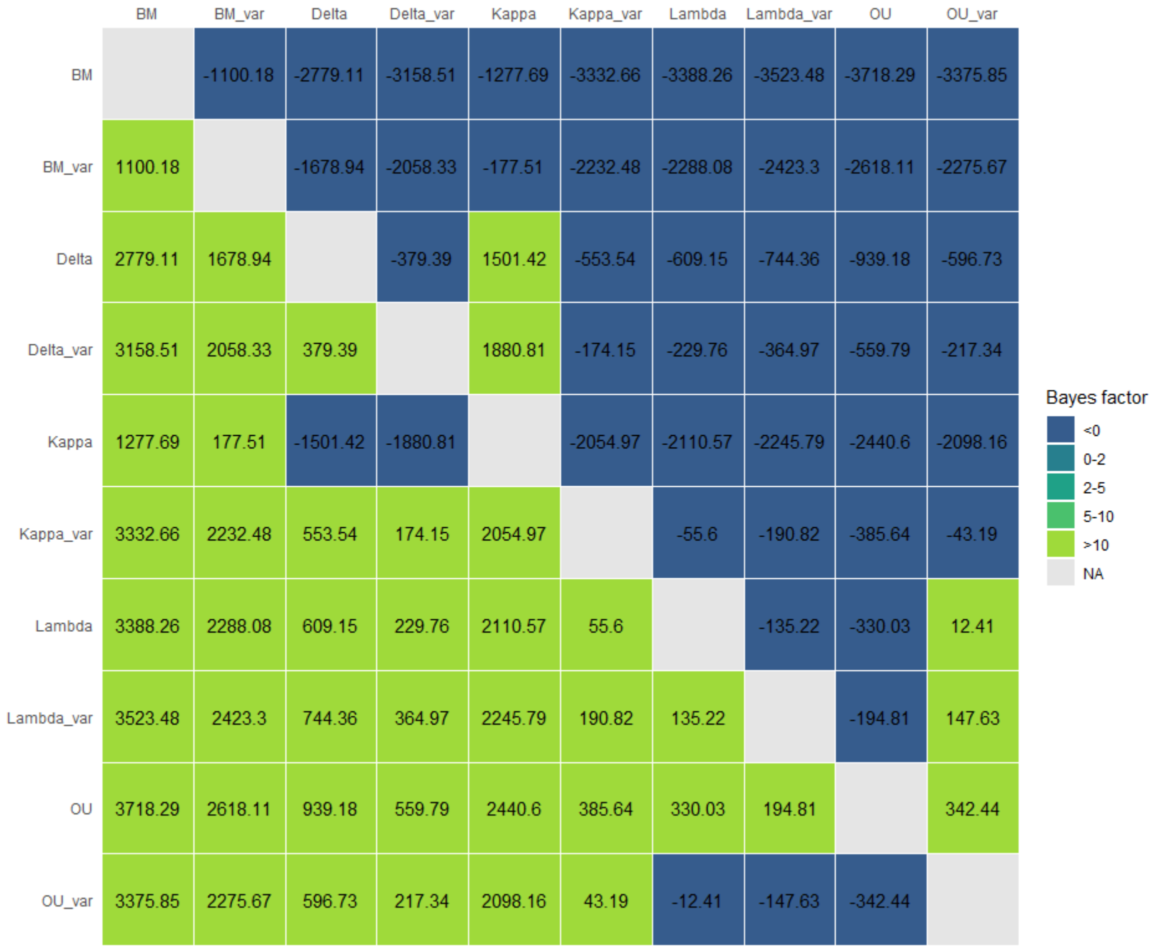


**Figure S3: Results of model comparison from BayesTraits analysis.**

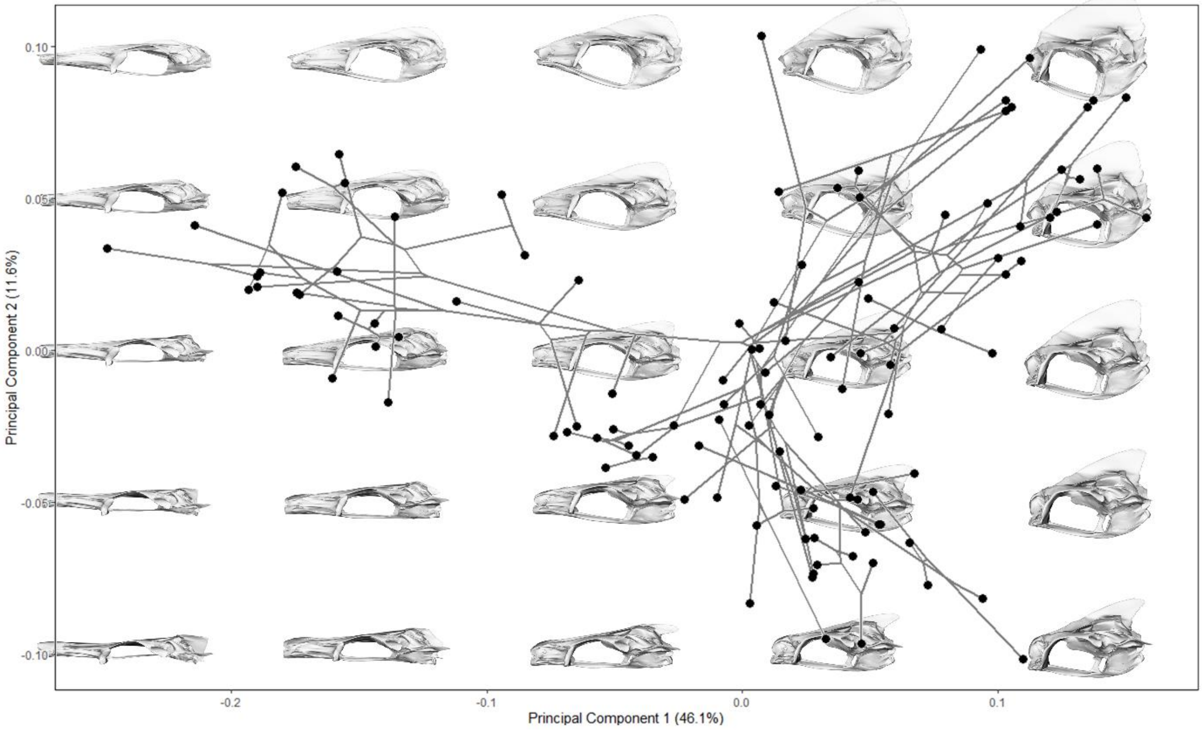


**Figure S4: Theoretical morphospace.** PCs 1 and 2 are shown with corresponding shape data, created from warping meshes to relevant coordinates.
